# Performance assessment of Respiratory Viral ELITe MGB® assay for the quantitative detection of influenza A/B and respiratory syncytial viruses

**DOI:** 10.1101/2020.05.14.097303

**Authors:** Antonio Piralla, Federica Giardina, Alice Fratini, Davide Sapia, Francesca Rovida, Fausto Baldanti

## Abstract

Influenza (Flu) and respiratory syncytial virus (RSV) are responsible for lower respiratory tract infections (LRTIs) associated with significant hospitalization among young children. In the present study, the performances of a triplex PCR assay detecting Flu A/B and RSV were compared with our in-house single-plex assays using 160 stored respiratory specimens previously tested using a panel of laboratory-developed real-time RT-PCR. Of them, 61 were positive for FluA, 41 for FluB, and 58 for RSV. All samples were retrospectively quantified with Respiratory Viral (RV) ELITe MGB® Panel (ELITechGroup Molecular Diagnostics, Puteaux, France) processed using ELITe InGenius® system. Overall, the total percentage agreement observed was 93.4% (57/61) for FluA, 92.7% (38/41) for FluB, and 86.2% (50/58) for RSV. A significant correlation of VL values was observed between the two methods for FluA and RSV (ρ= 0.91 and 0.84). This finding was supported by the strength of agreement between the two methods, as showed by the linear regression analysis (R2 =0.84 and 0.80). FluB viral load values measured by RV Panel were less significantly correlated (ρ= 0.77 and R2 =0.56). The bland-Altman analysis showed how 84.2% (48/57) of FluA and 86.0% of RSV (43/50) samples fell within ±1.0 Log10 variation from our laboratory results, while only 21.1% (8/38) of FluB results fell within this range. The great majority of FluB samples (29/30) outside range had values higher than +1.0 Log10 (median +2.1 Log10 range +1.0 to +3.5 Log10). In conclusion, RV ELITe MGB® Panel constitutes a valid and robust system for simultaneous detection and quantification of Flu A/B and RSV.

---

Influenza viruses type A and B (Flu A/B) and respiratory syncytial virus (RSV) are responsible for lower respiratory tract infections (LRTIs) associated with significant hospitalization among young children, elderly and immunocompromised patients (1–5). The incidence, morbidity, and mortality of Flu as compared to RSV varies from season to season (6). A rapid diagnosis allowing an appropriate decision regarding treatment and/or improved cohorting and isolation strategies to prevent transmission is a major concern on respiratory virus infections (7–9). In fact, in the last decade, the introduction of nucleic acid amplification tests (NAATs) have shortened turnaround time (TAT) and increased sensitivity for respiratory viruses (10). Furthermore, the multiplex RT-PCR approach is a validated strategy to detect a large number of respiratory viruses (8). Quantitative NAATs have been useful in terms of monitoring the reduction of viral load and thus the clinical efficacy of specific therapy (11,12). Different viral load levels have been associated with a higher risk of complications and severe disease in adults and children (13–15). In addition, the determination of viral load for different viruses in co-infections could be useful to distinguish which virus is the real pathogen and which the bystander (16). All these issues have to be interpreted in the context of available clinical and diagnostic information in order to improve clinical management. However, the use of quantitative NAATs in the diagnosis of respiratory viruses has largely been debated.

In the present study, the performances of a triplex-PCR assay detecting and quantifying Flu A/B and RSV were compared with our laboratory developed single-plex assays using positive stored clinical specimens.

## MATERIAL AND METHODS

### Study samples

A total of 160 respiratory samples, stored at −80 °C in a universal transport medium (UTM™, Copan Italia SpA, Brescia, Italy) and collected from December 2014 through April 2016 at the Molecular Virology Unit of the Fondazione IRCCS Policlinico San Matteo were included in this study. All samples were previously tested using a panel of laboratory-developed assays (LDA) real-time RT-PCR as previously described (9). Of them, 61 were positive for Flu A, 41 for Flu B and 58 RSV. Samples were categorized by viral load as high (>10^6^ RNA copies/ml), medium (10^4^-10^5^ RNA copies/ml) and low (10^2^-10^3^ RNA copies/ml). All samples were retrospectively quantified with Respiratory Viral ELITe MGB® Panel (ELITechGroup Molecular Diagnostics, Puteaux, France) processed using ELITe InGenius® system.

### Respiratory Viral ELITe MGB^®^ Panel

The archived respiratory samples were processed according to the manufacturer’s protocol on InGenius, a completely automated cassette based sample-to-results solution combining a universal extraction and independently controlled Real-time PCR thermal cycler (ELITechGroup Molecular Diagnostics, Puteaux, France). Briefly, 200 ul of respiratory were carefully transferred into a dedicated tube and loaded on the InGenius instrument for testing. Finally, the InGenius instrument was supplied with extraction/amplification Internal Control (IC), the RV ELITe MGB amplification Master mix, and extraction and amplification cassette consumables provided by the manufacturer (ELITechGroup Molecular Diagnostics, Puteaux, France). Results interpretation was performed according to the instruction manual of the RV ELITe MGB^®^ assay. Quantitative results expressed as log_10_ RNA copies/ml were measured comparing the cycle threshold (Ct) values obtained and interpolated with a standard curve (serial dilutions of DNA plasmid) for FluA, FluB and RSV.

### Statistical analysis

All viral RNA load (copies/ml) statistics were performed using log_10_ transformed viral load values. Quantitative variables were described as the mean and standard deviation, and/or median. Correlations between two quantitative variables were measured by the Spearman correlation test. The agreement between the assays was assessed with a Bland-Altman plot (17) and for graphical representation a ± 0.5 Log_10_ was considered an acceptable range of variability as also according to other publications (18). Descriptive statistics and linear regression lines were performed using Graph Pad Prism software (version 5.00.288). The correlation between the quantitative results was computed as the concordance correlation coefficient (CCC) of the measurements, according to Lin (19) using MedCalc® software (Version 9.4.2.0).

## RESULTS

A total of 160 respiratory samples with viral load ranging from 120 to 54574920 RNA copies/ml for FluA, from 180 to 31370040 RNA copies/ml for FluB and from 100 to 94513860 RNA copies/ml were analysed. Overall, the total percentage agreement observed was 93.4% (57/61) for FluA, 92.7% (38/41) for FluB and 86.2% (50/58) for RSV (Table 1). In detail, all FluA-(4/61) and FluB-positive (3/41) samples not detected by RV ELITe MGB® Panel belonged to low viral load group (10^2^-10^3^ RNA copies/ml), with viral load ranging from 270 to 900 RNA/copies ml for FluA and from 225 to 900 RNA/copies ml for FluB. Among 8 (13.8%) RSV-positive samples resulted negative by RV ELITe MGB® Panel, 1 (1.7%) had medium viral load (25650 RNA/copies ml) and 7 (12.1%) had low viral load ranging from 180 to 810 RNA/copies ml.

**TABLE 1.** Cross table of clinical performance study.

| Assay | Virus target | Samples viral load category | RV ELITe MGB® results |  |  | % agreement |
| --- | --- | --- | --- | --- | --- | --- |
|  |  |  | pos | neg | total |  |
| LDA | Flu A<br>(n=61) | high viral load<sup>a</sup> | 26 | 0 | 26 | 100.0% |
|  |  | medium viral load<sup>b</sup> | 17 | 0 | 17 | 100.0% |
|  |  | low viral load<sup>c</sup> | 14 | 4 | 18 | 77.7% |
|  |  | Total | 57 | 4 | 61 | 93.4% |
|  | Flu B<br>(n=41) | high viral load<sup>a</sup> | 11 | 0 | 11 | 100.0% |
|  |  | medium viral load<sup>b</sup> | 14 | 0 | 14 | 100.0% |
|  |  | low viral load<sup>c</sup> | 13 | 3 | 16 | 81.3% |
|  |  | Total | 38 | 3 | 41 | 92.7% |
|  | RSV<br>(n=58) | high viral load<sup>a</sup> | 13 | 0 | 13 | 100.0% |
|  |  | medium viral load<sup>b</sup> | 22 | 1 | 23 | 95.7% |
|  |  | low viral load<sup>c</sup> | 15 | 7 | 22 | 68.2% |
|  |  | Total | 50 | 8 | 58 | 86.2% |

LDA, laboratory developed assay; influenza A, Flu A; influenza B, Flu B; positive, pos; negative, neg
<sup>a</sup>>10<sup>6</sup> copies/ml
<sup>b</sup>10<sup>4</sup>-10<sup>5</sup> copies/ml
<sup>c</sup>10<sup>2</sup>-10<sup>3</sup> copies/ml

Positive samples were stratified based on viral load into three groups named high, medium and low (Figure 1). Viral loads were comparable in samples included in the high (p=0.11) and medium (p=0.84) group for FluA as well as for RSV (p=0.07 and p=0.74) (Fig. 1A and 1C). Conversely, a significantly difference of viral load was observed for FluA and RSV in low viral load group (Fig. 1A and 1C; p<0.001). For FluB samples, no difference in median viral load was observed in high group, while median viral load measured by RV ELITe MGB® Panel was greater in the medium and low groups (Fig. 2A, p<0.001).

**Figure 1.**
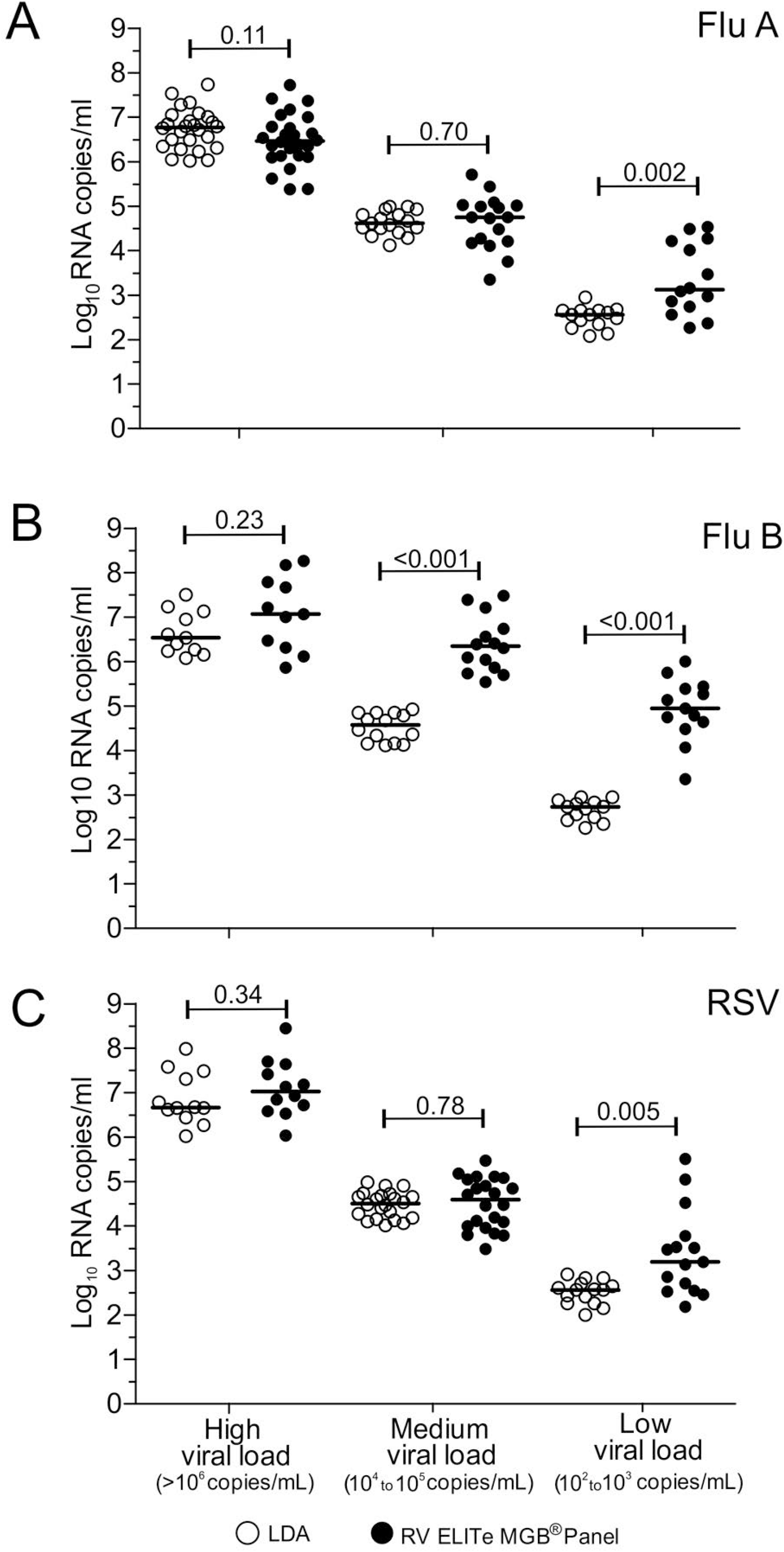
Comparison of the viral load measured with LDA (white circle) and RV ELITe MGB® Panel (black circle) for influenza A (A), influenza B (B) and RSV (C).

**Figure 2.**
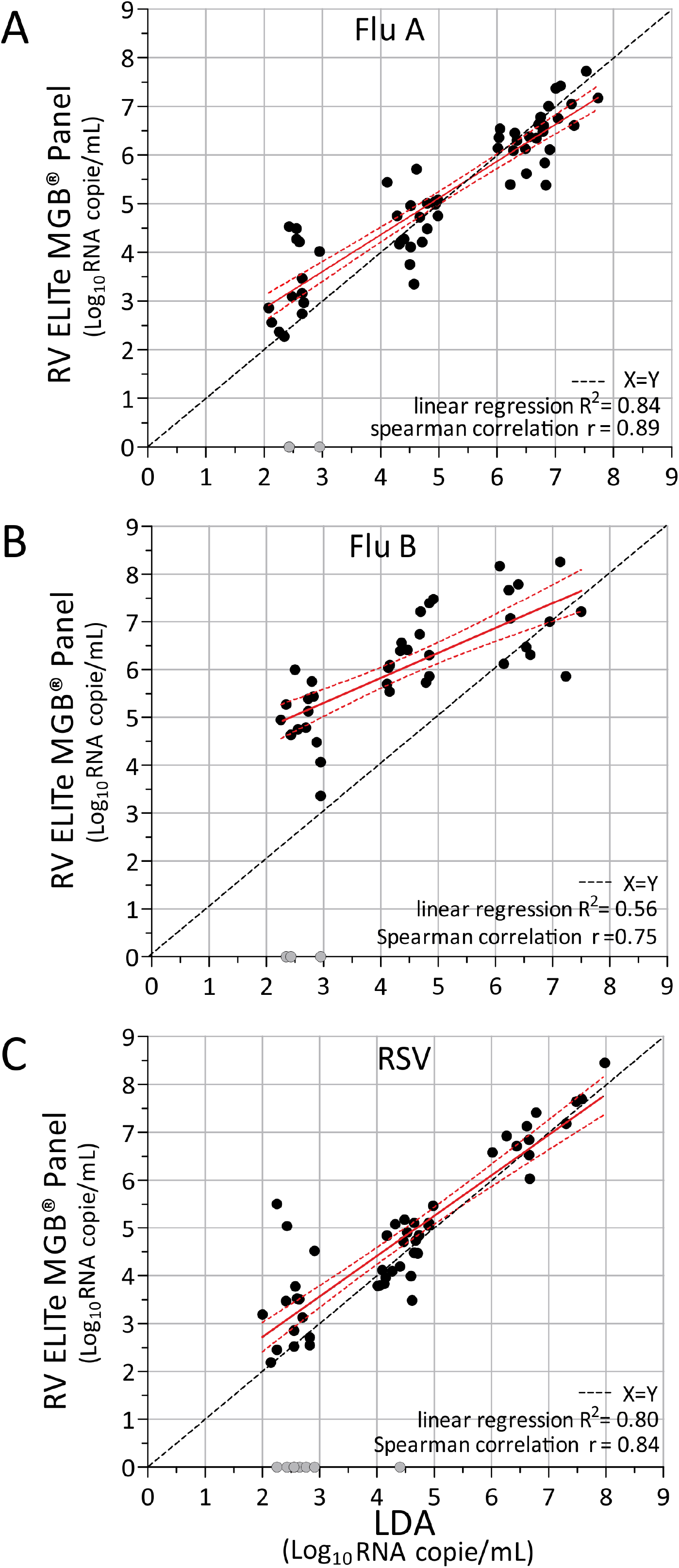
Linear regression analysis of log transformed viral load measured by LDA vs RV ELITe MGB® Panel assays for influenza A (A), influenza B (B) and RSV (C). Dashed red lines show the 95% confidence interval of the regression line (red line). Samples with undetectable results with RV ELITe MGB® Panel are reported with a grey circle.

A significant correlation was observed between the LDA and RV ELITe MGB® Panel for FluA and RSV assays (ρ= 0.91 and 0.84) also supported by the strength of agreement observed by the linear regression analysis (R^2^ =0.84 and 0.80) (Fig. 2A and 2C). In addition, the two assays showed good concordance, with a CCC of 0.86 (95% CI, 0.79 to 0.91) and 0.81 (95% CI, 0.70 to 0.88) for FluA and RSV assays, respectively (Table 2). FluB viral load values measured by RV ELITe MGB® Panel were less significant correlated to those quantified by LDA (ρ= 0.77 and R^2^ =0.56). This finding was also confirmed by the low concordance with a CCC of 0.36 (95% CI, 0.22 to 0.49).

**TABLE 2.** Results obtained from statistical analyses of two methods comparisons.

| Methods comparison | LDT vs RV ELITe MGB® Panel |  |  |
| --- | --- | --- | --- |
|  | Influenza A | Influenza B | RSV |
| Arithmetic mean (95% CI<sup>a</sup>) | -0.09 | -1.58 | -0.33 |
| Lower limit | -2.10 | -3.50 | -3.25 |
| Upper limit | +1.46 | +1.38 | +1.13 |
| SD | 0.73 | 1.08 | 0.74 |
| Concordance correlation coefficient | 0.86 | 0.36 | 0.81 |
| Person ρ (precision) | 0.89 | 0.75 | 0.84 |

Confidence interval, CI; standard deviation, SD

A plot of the differences between Log_10_ values obtained with LDA and results obtained by RV ELITe MGB® Panel assay was reported using a Bland-Altman analysis. Overall, the mean difference between two assays were −0.09 (± 1.96 SD, range −1.52 to +1.34) for FluA, −1.58 (± 1.96 SD, range −3.7 to +0.54) for FluB, and −0.32 (± 1.96 SD, range −1.77 to +1.13) for RSV. Assuming that differences within ± 0.5 log_10_ from LDA results for RV ELITe MGB® Panel assay is the acceptable range, Bland-Altman analysis showed how 64.9% (37/57) of FluA and 64.0% of RSV (32/50) samples fell within ±0.5 Log_10_ variation from LDA results, while only 15.8% (6/38) of FluB results fell within this range. Regarding values outside the range of acceptability (values >+0.5 log_10_ or <−0.5 log_10_ difference), among results of FluA samples, 10/57 (17.5%) had had values >+0.5 log_10_ difference (mean +0.88 Log_10_ range +0.51 to +1.46 Log_10_), while 10/57 (17.5%) had values <−0.5 log_10_ difference (mean −1.30 Log_10_ range −2.10 to −0.61 Log_10_). Among RSV samples, 3/50 (6.0%) had values >+0.5 log_10_ difference (mean +0.79 Log_10_ range +0.61 to +1.13 Log_10_) and 15/50 (30.0%) had values <−0.5 log_10_ difference (mean −1.14 Log_10_ range +3.25 to −0.51 Log_10_) (grey circle, Fig. 3A and 3C). Almost all (31/32) FluB samples outside the acceptability range had a viral load difference greater than −0.5 Log_10_ (median −1.99 Log_10_ range −0.80 to −3.5 Log_10_; Fig. 3B) as compared with our LDA. This means that overall RV ELITe MGB® Panel quantify 2 Log_10_ more than LDA.

**Figure 3.**
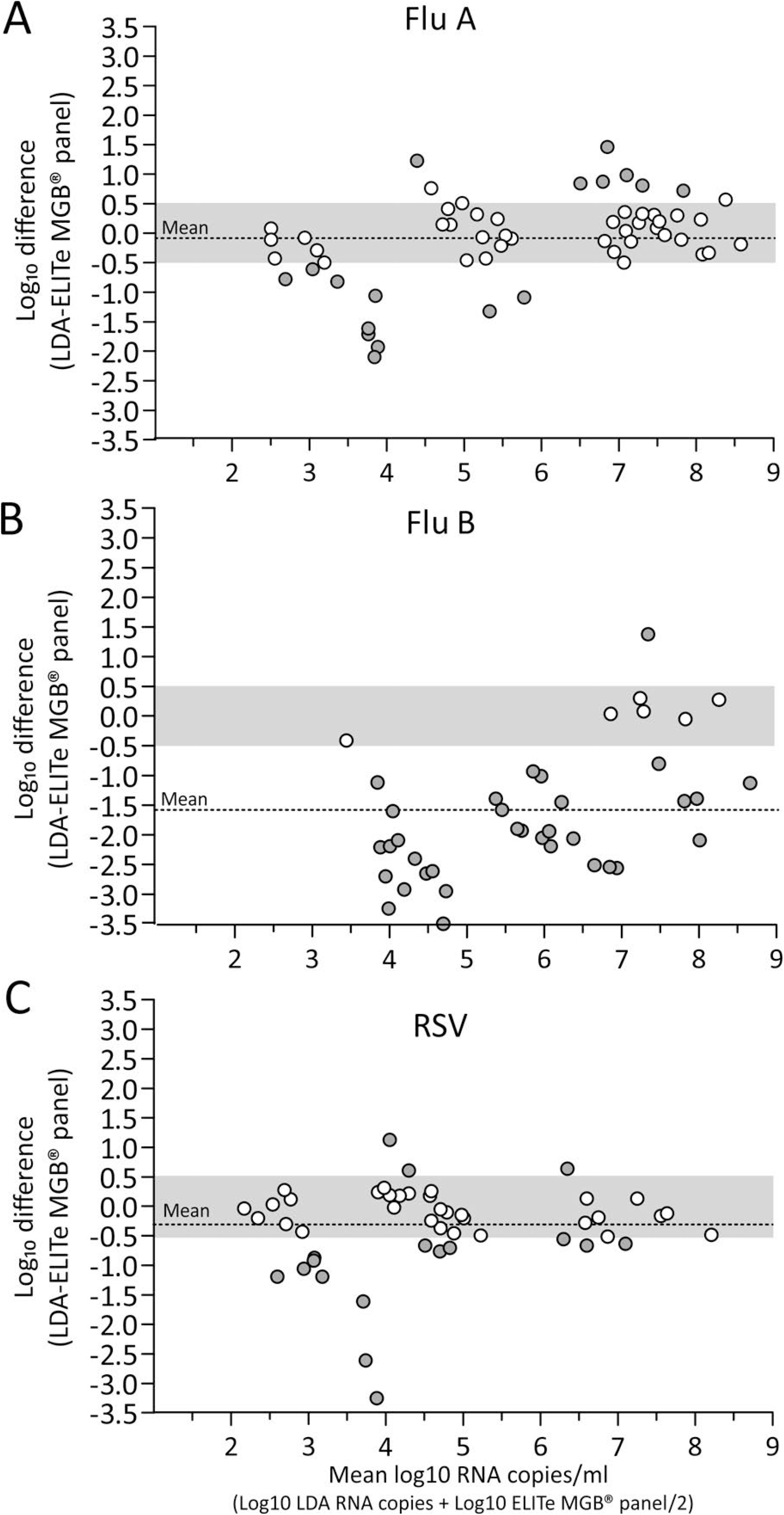
Bland Altman analysis was performed to compare the viral load (Log_10_ difference) measured by the two methods, LDA and RV ELITe MGB® Panel assay, for influenza A (A), influenza B (B) and RSV (C). The acceptability range (+0.5 to −0.5 Log_10_ difference) is shaded in light grey and mean value is reported with a dotted line. Data outside the acceptability range are reported with a grey circle.

## DISCUSSION

In the field of respiratory infections, a rapid and accurate diagnosis is needed for reducing unnecessary antibiotic usage, preventing transmission, and initiation of specific antiviral therapy (20, 21). In the past decades, the conventional diagnostics methods have been replaced by molecular assays also in the diagnosis of respiratory virus infections. Although, these assays have significantly reduced the turnaround time (TAT) to less than six hours, sometimes most of them resulted as complex to perform. In this perspective, it was of great introduction the newly designed diagnostic platform easy to handle with a further reduction of TAT. In the present study, the ELITe InGenius® system has been evaluated using the Respiratory Viral ELITe MGB® Panel in terms of performances including the semi-quantification of respiratory samples.

The overall agreement of the RV ELITe MGB® Panel compared to LDT was 93.4%, 92.7%, and 86.2% for influenza A, influenza B, and RSV, respectively. These findings are in keeping with the results of other rapid molecular assays when compared to LDT (22, 23). The main discordant results were observed in samples with low viral load (< 3 log10 RNA copies/ml). These results are commonly observed in comparison performed between multiplex syndromic PCR panels and single target LDT (24, 25). Linear regression showed good correlations between RV ELITe MGB® Panel and LDT for Flu A and RSV. Among Flu B samples, a greater viral load level with a median of 2 Log10 was observed using RV ELITe MGB® Panel.

RV ELITe MGB® Panel also provides a fully automated sample-to-result solution with a TAT of 2.5 hours for 12 samples but at this stage, only the panel does not include other respiratory pathogens. However, results of the present study encourage the availability of quantitative assays for respiratory virus detection but raise the question that also other respiratory viruses, such as rhinoviruses and parainfluenza viruses, could be included in a quantitative panel due to their increasing frequency of detection also in severe respiratory illness (26, 27).

Our study has a number of limitations. First, the RV ELITe MGB® Panel has been evaluated only in a series of previously tested-positive samples and therefore it could not be assessed an overall performance in terms of positive and negative predictive values. Our pilot study was mainly focused on the validation of the quantitative results obtained. It will be necessary, a more extended study should be performed, including also negative samples, in order to clarify the clinical impact of this sample-to-result solution within the laboratory workflow.

In conclusion, based on the data presented here, the robustness of quantification obtained by the RV ELITe MGB® Panel was demonstrated. Only a few samples with very low viral load have not been detected by the new direct RV ELITe MGB® Panel assay described herein is a powerful tool for rapid and simple molecular diagnosis of seasonal influenza as well as RSV.

## ACKNOWLEDGMENTS

The authors would like to thank Daniela Sartori for manuscript editing. Financial support and reagents were provided by ELITechGroup Molecular Diagnostics (Torino, Italy). The funding organization played no role in the collection, management, analysis, and interpretation of the data; or preparation, review, and approval of the manuscript.

